# Differential contributions of subthalamic beta rhythms and neural noise to Parkinson motor symptoms

**DOI:** 10.1101/312819

**Authors:** Stephanie Martin, Iñaki Iturrate, Ricardo Chavarriaga, Robert Leeb, Aleksander Sobolewski, Andrew M. Li, Julien Zaldivar, Iulia Peciu-Florianu, Etienne Pralong, Mayte Castro-Jiménez, David Benninger, François Vingerhoets, Robert T. Knight, Jocelyne Bloch, José del R. Millán

**Affiliations:** Defitech Chair in Brain-Machine Interface, Center for Neuroprosthetics, Ecole Polytechnique Fédérale de Lausanne, Switzerland; Helen Wills Neuroscience Institute, University of California, Berkeley, CA, USA; Li Ka Shing Faculty of Medicine, University of Hong Kong, Hong Kong, P.R. China; Department of Clinical Neurosciences (Neurology and Neurosurgery), University Hospital of Vaud (CHUV), Lausanne, Switzerland; Service de Neurochirurgie, Hôpital de Sion, Hôpital du Valais, Switzerland.; Department of Psychology, University of California, Berkeley, CA, USA

**Keywords:** Parkinson's disease, beta oscillatory activity, neural noise, power spectral density, biomarkers, closed-loop deep brain stimulation

## Abstract

**Background:** Excessive beta oscillatory activity in the subthalamic nucleus (STN) is linked to Parkinson’s disease and associated motor symptoms. However, the relationship between beta activity and motor symptoms has been inconsistent, which may influence the efficacy of closed-loop deep brain stimulation.

**Hypothesis:** We hypothesized that this variability is due to the degree of neural noise in STN recordings. Recent evidence has shown that neural noise is influenced by multiple factors, such as development, aging and disease, and could confound measures of beta activity. In this work, we propose a model that disentangles beta oscillatory activity and neural noise in the STN power spectrum.

**Methods:** We investigated the impact of neural noise on estimations of beta activity and motor symptoms from data recorded bilaterally from the subthalamic nuclei of thirteen Parkinsonian patients.

**Results:** Results showed that the relationship between beta oscillatory amplitude and motor symptoms (bradykinesia and rigidity) significantly improved when neural noise was removed from the estimation of beta activity.

**Conclusion:** These findings emphasize the importance of modeling neural components independently for understanding physiological processes associated with Parkinson’s disease, and identifying better biomarkers for characterizing symptom severity. Subsequently, we predict that our findings can have a direct application for closed-loop deep brain stimulation on Parkinson’s Disease.

## Introduction

Parkinson’s disease (PD) is a neurodegenerative disorder that is caused by a progressive loss of primarily dopaminergic neurons in the substantia nigra, leading toto various motor symptoms, such as bradykinesia, tremor and rigidity. Many studies have found that Parkinsonian patients have an excessive synchronization of beta oscillatory activity (13-30Hz) in local field potentials (LFPs) recorded from the subthalamic nucleus (STN) [1]. Enhanced beta activity is associated with bradykinesia and rigidity providing a biomarker of disease state. Both the dopamine precursor levodopa [2–5] and deep brain stimulation (DBS) [6–8] disrupt enhanced beta activity correlating with clinical motor improvement [9]. While it is clear that beta oscillatory activity is linked to motor impairments, its relationship with untreated clinical states (OFF-medications and OFF-stimulation) has been marked by inconsistent findings: While some physiological studies have shown a significant correlation with bradykinesia and rigidity[10–13], others have found no link [3–5].

Traditionally, beta oscillatory activity is not measured in an isolated manner, but rather measured together with the background noise activity of neurophysiological recordings, and it is reflected as a discrete *bump* in the power spectrum. This background noise, usually termed as neural noise activity, refers to random intrinsic electrical fluctuations within neuronal networks that display a 1/f^*n*^ power law behavior attributed to the low-pass frequency filtering property of the neural architecture and network mechanisms [14–17]. Recent works have shown that rich information is embedded in this neural noise activity, which thus carries physiological information. Indeed, it has already been demonstrated that the neural noise varies with several factors such as behavior, neural communication and cognitive impairments [18–20]. For instance, aging has been shown to be accompanied with a flatter power spectrum, attributed to an increase in spontaneous baseline neural activity observed both in intracranial and scalp EEG recordings [21]. These results suggest that neural noise activity could confound beta activity measurements, potentially explaining the large variability across studies.

In this study, we decomposed the LFP power spectrum as the sum of two independent neurophysiological elements –beta oscillatory activity and neural noise activity. We recorded STN LFPs bilaterally of thirteen PD patients, and modeled the beta activity with a Gaussian function and the neural noise with a power law function. We then evaluated the impact of noise when quantifying the relationship between beta oscillatory activity and motor symptoms. In addition, we assessed the stability of the neural noise and beta activity over time, in order to define the suitability of our approach for closed-loop deep brain stimulation [22].

## Materials and methods

### Subjects and experimental protocol

LFPs were recorded between 36 and 48 hours after STN DBS bilateral implantation in thirteen subjects using a 16-channel amplifier (gUsbAmp, g.tec, Austria) and sampled at 512 Hz. All patients volunteered and gave informed consent. Experimental protocol was approved by the local ethical committee (CER-VD, Commission Cantonale d’Ethique VD de la recherche sur l’être humain). Clinical characteristics of each patient are presented in Supplementary Table 1.

For each patient, LFPs were recorded bilaterally during six minutes in a resting state condition. Prior to recording, the stimulation device was turned off for at least ten minutes. Patients were instructed to relax and keep their eyes open during the recording. Patients were clinically assessed at the end of the recording by independent neurologists using the Unified Parkinson’s Disease Rating Scale (UPDRS) sub-scores for bradykinesia, rigidity and resting tremor symptoms for both upper limbs (items 20, 22 and 23).

### Surgical procedure

Long-acting dopaminergic medication was withdrawn 48 hours prior and short-acting medication was withdrawn 12-20 hours prior to off-medication pre-operative testing and to the DBS lead implantation procedure, always performed by the same surgeon. The awake surgical procedure stereotactically implanted a 3389 Medtronic electrode in the motor part of the STN. Direct surgical planning was performed with the Medtronic stealth station, based on a CT scan obtained with the CRW stereotactic frame, and fused with two 3 Tesla MRI sequences (MPRAGE with Gadolinium and space T2), obtained prior to surgery. After a first phase of microrecording, macrostimulation was performed under neurological clinical assessment. Adjustment of the electrode placement through a second track was eventually performed to optimize the clinical response to stimulation. The electrode location was checked during the surgery with the O-Arm (Medtronic). An externalized extension cable was connected to the distal part of the electrode and tunneled posterior to the skin incision to record the brain activity for a period of 3 days. Internalization of the electrode and connection of the electrode to an internal stimulator were performed at day 4 under general anesthesia.

### Power spectrum modeling

Six minutes of bipolar LFP activity were derived from two contacts, selected based on the electrodes chosen by the neurosurgeon for DBS stimulation. LFPs high-pass filtered at 1 Hz using a second order Butterworth filter. Power spectral densities (PSD) were then calculated using a Short-Time Fourier Transform, with a 0.25s Hamming window and 60% overlap and finally log powered.

We propose a model that describes the LFP spectrum and allows the extraction of two components of the PSD: the neural noise and the beta oscillatory activity. Similar approaches have been used to model the PSDs in EEG [23]. The different components of the model are illustrated in Fig. 1A, in which the LFP PSDs are modeled as:

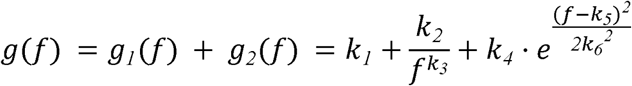

where g(*f*)is the modeled power spectrum at frequency *f*. The function *g*_1_(*f*) models the neural LFP noise function at frequency *f* with a power law function; *k*_1_, *k*_2_, *k*_3_, determine its offset, rotational offset and power, respectively. The offset was added to the equation so as to shift the axis of overall rotation (intersection frequency) that would be fixed at 1Hz if only the rotational offset was taken into account. The function *g*_2_(*f*) models the beta activity function at frequency *f* with a Gaussian probability density function, with amplitude *k*_4_, mean frequency *k*_5_, and standard deviation *k*_6_. *k*_1–6_ parameters were estimated from the entire 6-min recording by an interior-point optimization algorithm, with the objective function defined as the root mean square error between the modeled and actual PSDs:

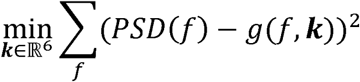

where frequencies *f* ranged between 10-90 Hz. Lower and upper bounds on the design variables were defined based on a priori knowledge and the shape of typical power spectral densities [23]. Initial values of parameters were set to the central value over the range defined by the lower and upper bounds (see Supplementary Table 2).

**Figure 1:**
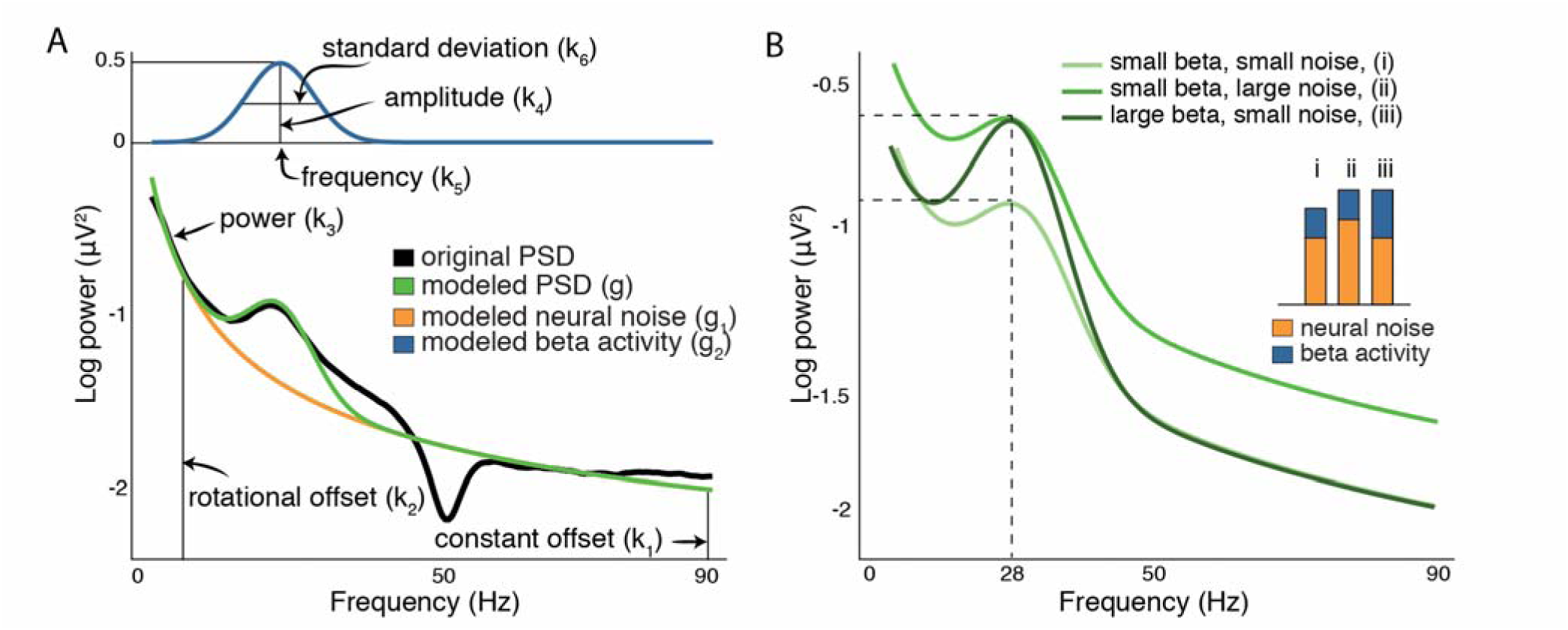
LFP modelization. **A.** The model assumes two components: the neural noise activity and the beta activity. **B.**Three examples of power spectrum with different levels of beta oscillatory activity and neural noise activity. The bar graph represents the level of noise activity and beta activity.

Fig. 1B exemplifies how the neural noise level might affect the quantification of beta oscillatory activity. The light green curve shows a small beta amplitude and a low level of neural noise evaluated at the peak amplitude (28Hz). Alternatively, the neutral green curve has the same beta amplitude as the light green curve, but the level of neural noise is larger, resulting in an overall larger total amplitude. Finally, the dark green curve has the same overall amplitude as the neutral green curve, but the beta amplitude is actually much larger. These examples emphasize the need to model these physiological processes independently.

Emerging evidence has also shown the existence of two distinct beta sub-bands carrying pathological information [24]: the low-beta (13–20 Hz) and the high-beta (20–35 Hz) bands. As such, we also evaluated the feasibility to model the PSDs with two beta bumps. To this end, the PSD was modeled as the sum of two Gaussian function and a power law function, and therefore estimated nine parameters instead of six parameters in the case of a single beta bump.

### Statistical analysis

To validate our model, we first evaluated the goodness-of-fit between the modeled and actual PSDs using the coefficient of determination (*R*^2^). We then quantified the relationship between the model parameters *k*_1–6_ and the clinical scores using Spearman’s correlation coefficient (rho), pooling patients and hemispheres (N=26). The significance level was computed using permutation statistics and corrected for multiple comparisons using false discovery rate (FDR). We then compared s differences in correlation with clinical scores, when the beta oscillatory activity was estimated with and without neural noise. The estimated beta activity without noise corresponded to the amplitude of the Gaussian function (*k*_4_). The beta activity estimated with noise corresponded to the amplitude of the Gaussian function (*k_4_*) plus the value of the modeled neural noise function evaluated at the center frequency of the Gaussian function, *g*_1_(*k*_5_). We assessed for significant differences between correlation values with and without neural noise using Hotelling’s t statistic. This statistical test accounts for the dependence of the two correlations on the same group, i.e. both correlations are relative to the same motor symptoms. To investigate the relationship between the neural noise activity and motor symptoms, we also computed Spearman’s correlation between the modeled neural noise function and clinical scores at each frequency bin (1–200 Hz). Similarly, we measured the correlation between the neural noise activity and age.

## Results

### Modeling of the power spectrum

Goodness-of-fit (*R*^2^) between the modeled and actual PSDs ranged between 0.95-0.99, and thus our mathematical model accurately fitted the power spectrum. Similarly, goodness-of-fit obtained when modeling two beta bumps ranged between 0.94-0.99. Although this result suggests that our approach can be used to investigate low and high beta bumps independently, the adjusted *R*^2^– which takes into account the number of parameters as a regularization method– was not significantly different between the modeled PSDs with one or two beta bumps (Wilcoxon signed rank; *P=0.16*), suggesting an overfitting effect when modeling two bumps. Thus we further investigated the role of beta activity using only a single beta bump.

### Correlation between model parameters and motor symptoms

In order to identify how the LFP power relate to motor symptoms, we investigated the relationship between the different parameters of the modeled power spectrum and the UPDRS clinical scores. The fitted amplitude of the beta activity (*k_4_*) correlated with both bradykinesia (rho=0.55, *P*=0.006) and rigidity (rho=0.69, *P*=0.000), but not with tremor (rho=−0.07, *P*=0.743; Fig. 2). In addition, the beta amplitude correlated with bradykinesia plus rigidity (rho=0.68, *P*=0.000) and all three clinical scores summed together (rho=0.50, *P*=0.011). No other parameters presented significant correlations (*P*>0.1; Supplementary Table 3). These findings are in agreement with previous electrophysiological studies showing that pathological beta activity was associated with bradykinesia and rigidity, but not tremor [1].

**Figure 2:**
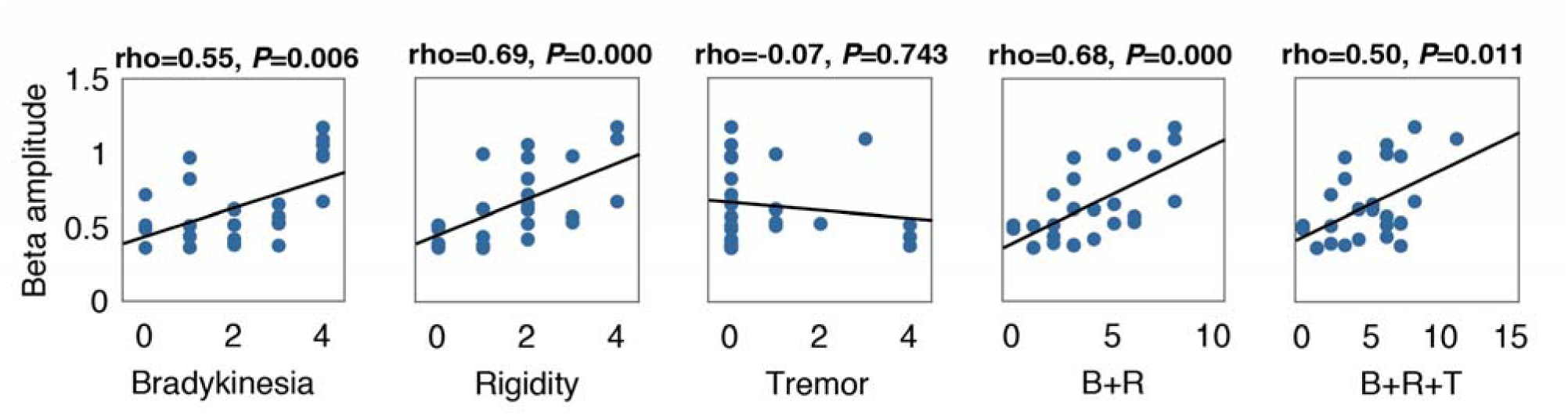
Relationship between modeled beta activity and motor symptoms. Correlation between beta amplitude estimated from the model (*k*_4_) and motor symptoms. Each panel shows the estimated beta amplitude value for each patient and hemisphere (N=26), together with least-squares regression line (black line), and the Spearman’s correlation and associated p-value on top (FDR correction). B=bradykinesia, R=rigidity and T=tremor.

### Impact of neural noise on the estimation of beta oscillatory activity

As neural noise is known to vary with factors such as behavior, neural communication and cognitive impairments [18,19,21], we assessed the influence of neural noise when estimating the relationship between beta oscillatory activity and motor symptoms. Fig. 3 shows the correlation between the clinical scores and the modeled beta activity with and without neural noise. The correlation without neural noise was increased compared to the estimation with neural noise for bradykinesia, rigidity, bradykinesia plus rigidity, and all three clinical scores summed together (*P*<0.05, Hotelling’s t-test), but not for tremor (*P*=0.316, Hotelling’s t-test). The correlations between motor symptoms and beta activity with noise did not reach significance: bradykinesia (rho=0.34, *P*=0.149), rigidity (rho=0.39, *P*=0.129), bradykinesia plus rigidity (rho=0.28, *P*=0.129) and all three summed together(rho=−0.13, *P*=0.534). The improvements with noise elimination ranged between 60-80% (rho_without_-rho_with_)/rho_with_), and demonstrate that the estimation of beta oscillatory activity correlates better with motor symptoms when it is isolated from the neural noise component.

**Figure 3:**
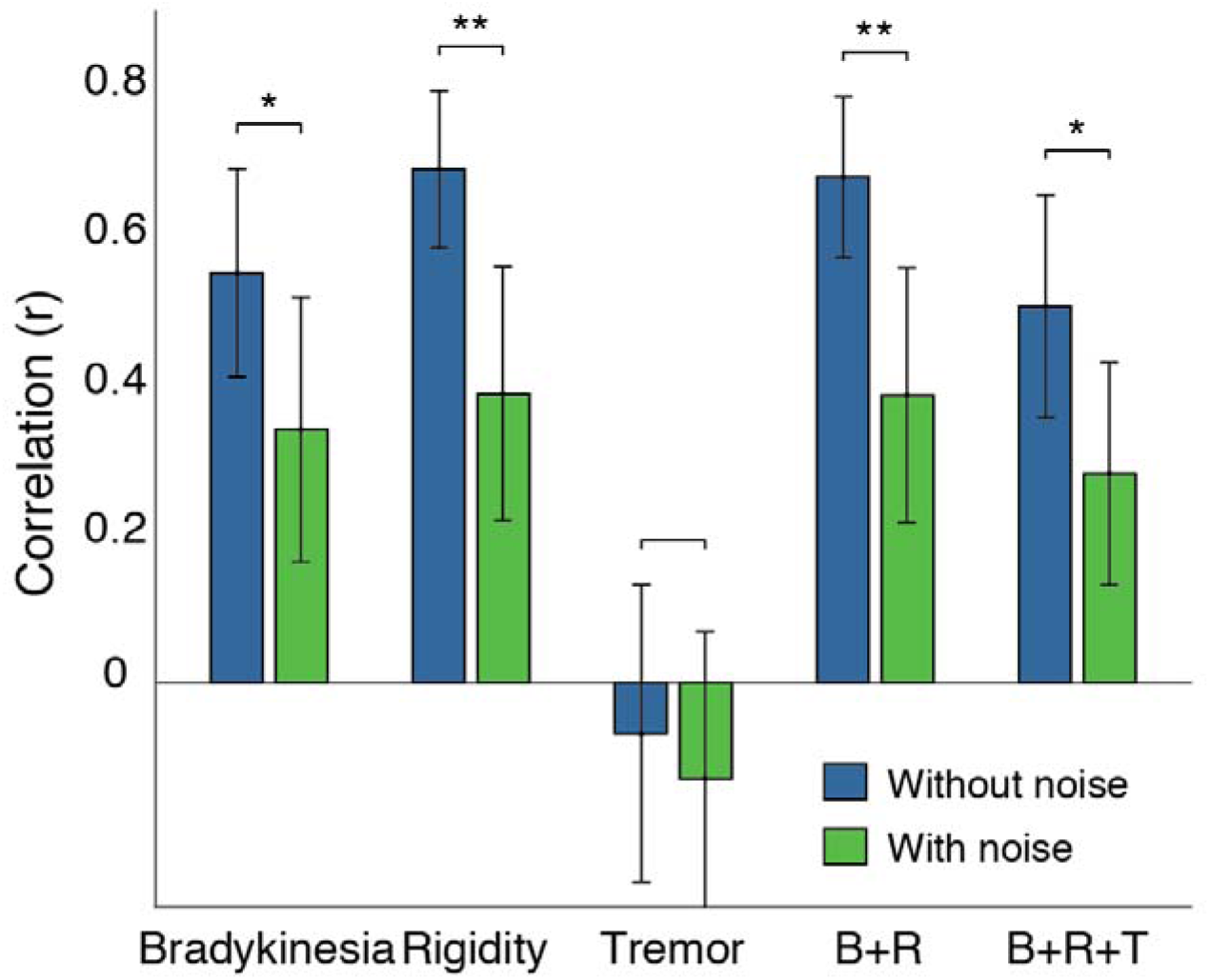
Impact of neural noise on beta activity estimations. Spearman’s correlation coefficient between beta oscillatory activity and motor symptoms, with and without neural noise. Error bars denote standard deviation marked (*P<0.05, **P<0.01, Hotelling’s t-test, FDR correction). B=bradykinesia, R=rigidity and T=tremor.

We also evaluated the within-subject non-stationarity of neural noise, measured as its variability throughout the time. The rationale is that a higher variability of neural noise would imply a noisier, less reliable estimation of any ongoing pathological biomarker that includes neural noise, beta oscillations in particular. This is particularly relevant for implementing closed-loop DBS paradigms, where the stimulation is controlled based on the estimated beta power [25,26]. We thus evaluated the variability of the estimated beta activity and neural noise over time by modeling,for each hemisphere and patient, the PSDs with a sliding window of ten seconds and no overlap, and computed the standard deviation across the time windows. Results showed that the variability throughout time of beta activity was significantly smaller than that of the neural noise (variability for beta activity (mean ± std across subjects) = 0.026 ± 0.028; neural noise = 0. 0310 ± 0.027; Wilcoxon signed rank test; *P*<10^−4^). Expectedly, the variability of the original PSD beta computation without our model – as usually done in typical closed-loop scenarios [25] – was even higher than the neural noise (mean ± std across subjects = 0.057 ± 0.055). This significantly larger standard deviation of neural noise with respect to beta activity suggests that modeling beta activity and neural noise independently might be beneficial for closed-loop DBS.

### Age-related changes in neural noise activity

Recent studies have shown that rich information is embedded in the neural noise activity, and the shape of the PSD has been shown to correlate with age and cognitive decline [21]. As such, we assessed the relationships between the neural noise activity in the STN and age, as well as motor symptoms. For this, we estimated the neural noise activity with the parameters *k*_1–3_, (offset, rotational offset and power of the neural noise, respectively function *g_1_*), and correlated the power at each frequency bin with motor symptoms. Fig. 4 shows the correlation between the modeled neural noise activity *g_1_* and each of the motor symptoms. Results showed no significant correlation between the neural noise activity and motor symptoms (*P*>0.05). Conversely, we found a significant negative correlation between age and neural noise activity for frequencies ranging between 7-21 Hz (*P*<0.05), and a positive correlation for frequencies ranging between 150-200 Hz, peaking at 163 Hz (*P*<0.05), in line with previous works [20]. These results strongly indicate the physiological nature of the neural noise (from its correlation with age), and its independence from other physiological signal and pathological symptoms (from the lack of correlation with PD motor symptoms).

**Figure 4:**
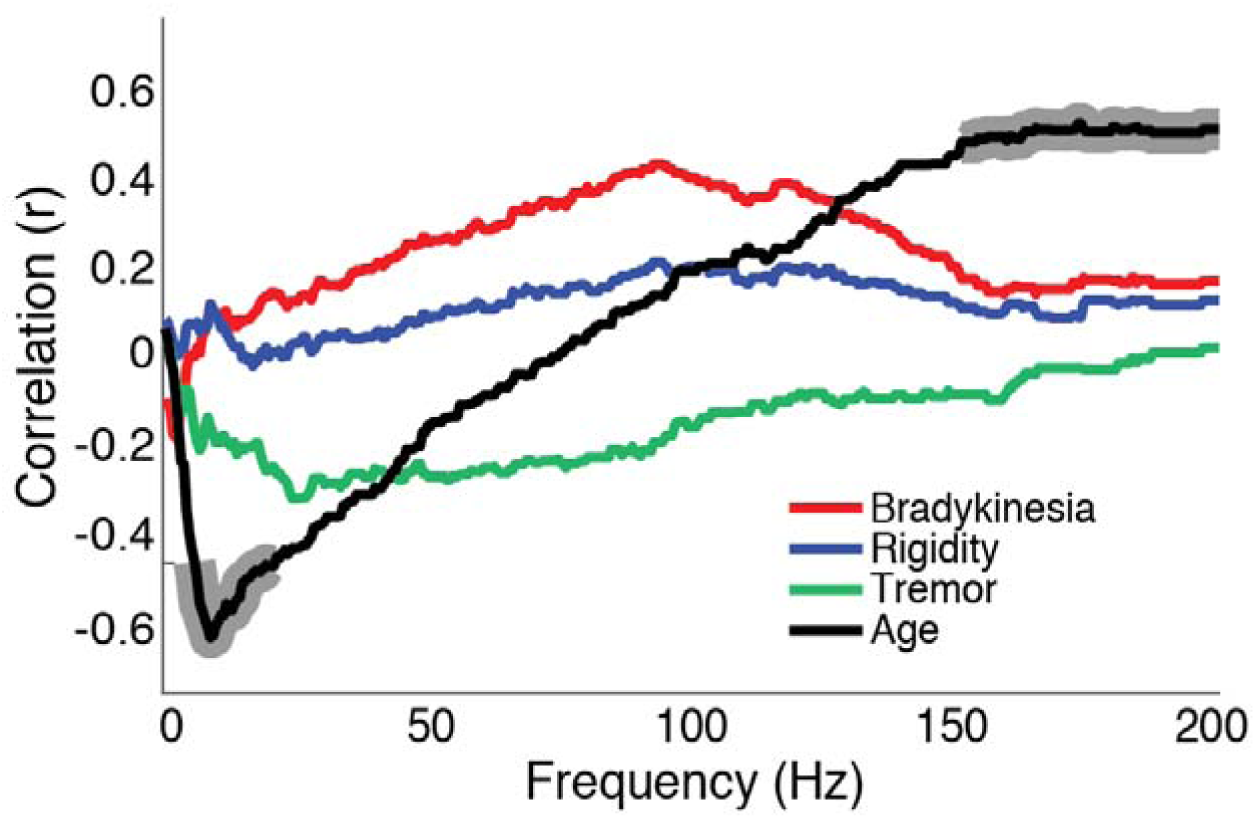
Correlation of neural noise with motor symptoms and age. Spearman’s correlation between the neural noise activity and each motor symptom and age, at each frequency bin. Significant correlations are marked by shadowed regions (*P*<0.05, FDR correction).

## Discussion

In this study, we modeled the LFP power spectrum in the STN of PD patients as the combination of two distinct physiological components: beta oscillatory activity and neural noise activity, modeled with a Gaussian function and a 1/*f*^*n*^ power law function. The modeled beta amplitude correlated with bradykinesia and rigidity, but not tremor, thereby reinforcing the concept that tremor is pathophysiologically distinct from bradykinesia and rigidity. Critically, the correlation between the estimated beta oscillatory amplitude and bradykinesia and rigidity was significantly enhanced when the neural noise activity was removed from the beta estimation. Finally, neural noise correlated with age, but not with motor symptoms, highlighting the differential roles of oscillatory activity and broadband activity in PD. Our model provides a tool to disentangle physiological processes associated with STN-LFPs, and allows further investigations on the neurobiological origin of the neural noise and its implications in PD and other neurological disorders.

Despite advances in the understanding of the pathophysiology underlying Parkinson’s disease, the causal role of beta activity remains unclear, as evidence has shown contradictory findings in untreated resting PD states. Indeed, studies that found a significant correlation have used various forms of power normalization or more complex measures, such as entropy [27]. Although these methods have implicitly tackled the variability induced by neural noise, they have not explicitly addressed the impact of this neural noise, nor suggested its differential role in motor state estimation. In this study, we evaluated for the first time the impact of the neural noise on the estimation of beta activity. By isolating the beta activity from the noise, we were able to obtain an increase in the correlation coefficients between motor symptoms and LFP beta activity by 60-80%. These results suggest that neural noise, while not correlating with motor symptoms, still confounded the estimations of beta oscillatory activity.

Our approach also allows identifying whether the changes in beta activity are caused by an actual drop in the beta oscillatory amplitude or rather by changes in the overall background level of activity. In this study, neural noise and age correlated negatively with low frequencies (7-21Hz) and positively with high frequencies (150-200Hz). These findings emphasize its physiological origins, and are in line with previous works: arecent study demonstrated that ECoG and EEG electrophysiological neural noise correlates with age and cognitive impairments [21], and it has been associated with an increase in baseline neural spiking activity [28]. It has been hypothesized that increases in the neural noise are a consequence of pathological decoupling between population spiking activity and low-frequency oscillatory neural field [18], and is associated with an imbalance between synaptic excitation and inhibition [29]. Further research is needed to define how background neural activity and oscillatory components interact in PD and how it can affect neural communication in the basal-ganglia-thalamocortical network.

The model proposed here provides promising new avenues for computing better biomarkers associated with PD symptoms. Indeed, we showed that the model can be customized depending on the shape of the PSDs, e.g. modeling two bumps for low and high beta frequency bands. As such, this modeling approach allows capturing and isolating independent neural processes. In addition, it is fully automatic, data-driven and personalized to patients, without needing any human intervention.

Our two-components modeling technique has direct applications for closed-loop DBS. Despite the benefits of continuous DBS, several studies have reported different side-effects, affecting mood, behavior, speech and personality [30]. One way to minimize the associated side-effects of DBS is to reduce the stimulation time using closed-loop systems, limiting the appearance of motor symptoms measured by specific biomarkers. As such, there is a strong interest in finding stable and reliable biomarkers for closed-loop DBS. Currently, beta oscillatory power has been used as a DBS trigger to reduce the stimulation time and improve motor performances compared to continuous stimulation [22,25,26]. Previous works have already suggested that neural noise does vary with time, and could do so at the sub-second level [18]. In this work, we have further shown that this variability exists, and that is significantly higher than the variability of the beta activity. As such, the proposed modeling approach could lead to more reliable quantifications of beta activity as biomarker for future closed-loop DBS applications [31].

## Acknowledgements

We thank Michael Pereira and Elvira Pirondini for their comments on the manuscript and help on the recordings. I.I., R.C., R.L., A.S., J.M. were supported by the Swiss National Centres of Competence in Research (NCCR) Robotics. I.I. also acknowledges support from the ‘EPFL Fellows’ fellowship program co-funded by Marie Curie, FP7 Grant agreement no. 291771. RTK is supported by NINDS R37NS21135. DB is supported by the Foundation Parkinson Switzerland, the Swiss Dystonia Society and the Swiss Accident Insurance Fund, and holds the Baasch-Medicus Prize.

## References

[1] C. Hammond, H. Bergman, P. Brown, Pathological synchronization in Parkinson’s disease: networks, models and treatments, Trends Neurosci. 30 (2007) 357–364. doi:10.1016/j.tins.2007.05.004.

[2] A.A. Kühn, A. Kupsch, G.-H. Schneider, P. Brown, Reduction in subthalamic 8-35 Hz oscillatory activity correlates with clinical improvement in Parkinson’s disease: STN activity and motor improvement, Eur. J. Neurosci. 23 (2006) 1956–1960. doi:10.1111/j.1460-9568.2006.04717.x.

[3] A.A. Kühn, A. Tsui, T. Aziz, N. Ray, C. Brücke, A. Kupsch, G.-H. Schneider, P. Brown, Pathological synchronisation in the subthalamic nucleus of patients with Parkinson’s disease relates to both bradykinesia and rigidity, Exp. Neurol. 215 (2009) 380–387. doi:10.1016/j.expneurol.2008.11.008.

[4] M. Weinberger, N. Mahant, W.D. Hutchison, A.M. Lozano, E. Moro, M. Hodaie, A.E. Lang, J.O. Dostrovsky, Beta Oscillatory Activity in the Subthalamic Nucleus and Its Relation to Dopaminergic Response in Parkinson’s Disease, J. Neurophysiol. 96 (2006) 3248–3256. doi:10.1152/jn.00697.2006.

[5] N.J. Ray, N. Jenkinson, S. Wang, P. Holland, J.S. Brittain, C. Joint, J.F. Stein, T. Aziz, Local field potential beta activity in the subthalamic nucleus of patients with Parkinson’s disease is associated with improvements in bradykinesia after dopamine and deep brain stimulation, Exp. Neurol. 213 (2008) 108–113. doi:10.1016/j.expneurol.2008.05.008.

[6] P. Brown, A. Oliviero, P. Mazzone, A. Insola, P. Tonali, V. Di Lazzaro, Dopamine dependency of oscillations between subthalamic nucleus and pallidum in Parkinson’s disease, J. Neurosci. Off. J. Soc. Neurosci. 21 (2001) 1033–1038.

[7] A. Eusebio, W. Thevathasan, L. Doyle Gaynor, A. Pogosyan, E. Bye, T. Foltynie, L. Zrinzo, K. Ashkan, T. Aziz, P. Brown, Deep brain stimulation can suppress pathological synchronisation in parkinsonian patients, J. Neurol. Neurosurg. Psychiatry. 82 (2011) 569–573. doi:10.1136/jnnp.2010.217489.

[8] D. Whitmer, C. de Solages, B. Hill, H. Yu, J.M. Henderson, H. Bronte-Stewart, High frequency deep brain stimulation attenuates subthalamic and cortical rhythms in Parkinson’s disease, Front. Hum. Neurosci. 6 (2012) 155. doi:10.3389/fnhum.2012.00155.

[9] P. Hickey, M. Stacy, Deep Brain Stimulation: A Paradigm Shifting Approach to Treat Parkinson’s Disease, Front. Neurosci. 10 (2016). doi:10.3389/fnins.2016.00173.

[10] J. Lopez-Azcarate, M. Tainta, M.C. Rodriguez-Oroz, M. Valencia, R. Gonzalez, J. Guridi, J. Iriarte, J.A. Obeso, J. Artieda, M. Alegre, Coupling between Beta and High-Frequency Activity in the Human Subthalamic Nucleus May Be a Pathophysiological Mechanism in Parkinson’s Disease, J. Neurosci. 30 (2010) 6667–6677. doi:10.1523/JNEUROSCI.5459-09.2010.

[11] A. Pogosyan, F. Yoshida, C.C. Chen, I. Martinez-Torres, T. Foltynie, P. Limousin, L. Zrinzo, M.I. Hariz, P. Brown, Parkinsonian impairment correlates with spatially extensive subthalamic oscillatory synchronization, Neuroscience. 171 (2010) 245–257. doi:10.1016/j.neuroscience.2010.08.068.

[12] T.E. Özkurt, M. Butz, M. Homburger, S. Elben, J. Vesper, L. Wojtecki, A. Schnitzler, High frequency oscillations in the subthalamic nucleus: A neurophysiological marker of the motor state in Parkinson’s disease, Exp. Neurol. 229 (2011) 324–331. doi:10.1016/j.expneurol.2011.02.015.

[13] W.-J. Neumann, K. Degen, G.-H. Schneider, C. Brücke, J. Huebl, P. Brown, A.A. Kühn, Subthalamic synchronized oscillatory activity correlates with motor impairment in patients with Parkinson’s disease: Correlation of Subthalamic Oscillations and PD Symptoms, Mov. Disord. 31 (2016) 1748–1751. doi:10.1002/mds.26759.

[14] C. Gold, D.A. Henze, C. Koch, G. Buzsáki, On the origin of the extracellular action potential waveform: A modeling study, J. Neurophysiol. 95 (2006) 3113–3128. doi:10.1152/jn.00979.2005.

[15] K.H. Pettersen, E. Hagen, G.T. Einevoll, Estimation of population firing rates and current source densities from laminar electrode recordings, J. Comput. Neurosci. 24 (2008) 291–313. doi:10.1007/s10827-007-0056-4.

[16] H. Lindén, K.H. Pettersen, G.T. Einevoll, Intrinsic dendritic filtering gives low-pass power spectra of local field potentials, J. Comput. Neurosci. 29 (2010) 423–444. doi:10.1007/s10827-010-0245-4.

[17] G. Buzsáki, X.-J. Wang, Mechanisms of gamma oscillations, Annu. Rev. Neurosci. 35 (2012) 203–225. doi:10.1146/annurev-neuro-062111-150444.

[18] W.J. Freeman, J. Zhai, Simulated power spectral density (PSD) of background electrocorticogram (ECoG), Cogn. Neurodyn. 3 (2009) 97–103. doi:10.1007/s11571-008-9064-y.

[19] E. Podvalny, N. Noy, M. Harel, S. Bickel, G. Chechik, C.E. Schroeder, A.D. Mehta, M. Tsodyks, R. Malach, A unifying principle underlying the extracellular field potential spectral responses in the human cortex, J. Neurophysiol. 114 (2015) 505–519. doi:10.1152/jn.00943.2014.

[20] B. Voytek, R.T. Knight, Dynamic Network Communication as a Unifying Neural Basis for Cognition, Development, Aging, and Disease, Biol. Psychiatry. 77 (2015) 1089–1097. doi:10.1016/j.biopsych.2015.04.016.

[21] B. Voytek, M.A. Kramer, J. Case, K.Q. Lepage, Z.R. Tempesta, R.T. Knight, A. Gazzaley, Age-Related Changes in 1/f Neural Electrophysiological Noise, J. Neurosci. 35 (2015) 13257–13265. doi:10.1523/JNEUROSCI.2332-14.2015.

[22] G. Tinkhauser, A. Pogosyan, S. Little, M. Beudel, D.M. Herz, H. Tan, P. Brown, The modulatory effect of adaptive deep brain stimulation on beta bursts in Parkinson’s disease, Brain J. Neurol. 140 (2017) 1053–1067. doi:10.1093/brain/awx010.

[23] B. Blankertz, C. Sannelli, S. Halder, E.M. Hammer, A. Kübler, K.-R. Müller, G. Curio, T. Dickhaus, Neurophysiological predictor of SMR-based BCI performance, NeuroImage. 51 (2010) 1303–1309. doi:10.1016/j.neuroimage.2010.03.022.

[24] A. Priori, G. Foffani, A. Pesenti, F. Tamma, A.M. Bianchi, M. Pellegrini, M. Locatelli, K.A. Moxon, R.M. Villani, Rhythm-specific pharmacological modulation of subthalamic activity in Parkinson’s disease, Exp. Neurol. 189 (2004) 369–379. doi:10.1016/j.expneurol.2004.06.001.

[25] S. Little, A. Pogosyan, S. Neal, B. Zavala, L. Zrinzo, M. Hariz, T. Foltynie, P. Limousin, K. Ashkan, J. FitzGerald, A.L. Green, T.Z. Aziz, P. Brown, Adaptive deep brain stimulation in advanced Parkinson disease: Adaptive DBS in PD, Ann. Neurol. 74 (2013) 449–457. doi:10.1002/ana.23951.

[26] S. Little, M. Beudel, L. Zrinzo, T. Foltynie, P. Limousin, M. Hariz, S. Neal, B. Cheeran, H. Cagnan, J. Gratwicke, T.Z. Aziz, A. Pogosyan, P. Brown, Bilateral adaptive deep brain stimulation is effective in Parkinson’s disease, J. Neurol. Neurosurg. Psychiatry. 87 (2016) 717–721. doi:10.1136/jnnp-2015-310972.

[27] C.C. Chen, Y.T. Hsu, H.L. Chan, S.M. Chiou, P.H. Tu, S.T. Lee, C.H. Tsai, C.S. Lu, P. Brown, Complexity of subthalamic 13-35Hz oscillatory activity directly correlates with clinical impairment in patients with Parkinson’s disease, Exp. Neurol. 224 (2010) 234–240. doi:10.1016/j.expneurol.2010.03.015.

[28] S.L. Hong, G.V. Rebec, A new perspective on behavioral inconsistency and neural noise in aging: compensatory speeding of neural communication, Front. Aging Neurosci. 4 (2012). doi:10.3389/fnagi.2012.00027.

[29] R. Gao, E.J. Peterson, B. Voytek, Inferring synaptic excitation/inhibition balance from field potentials, NeuroImage. 158 (2017) 70–78. doi:10.1016/j.neuroimage.2017.06.078.

[30] D. Cyron, Mental Side Effects of Deep Brain Stimulation (DBS) for Movement Disorders: The Futility of Denial, Front. Integr. Neurosci. 10 (2016). doi:10.3389/fnint.2016.00017.

[31] I. Iturrate, M. Pereira, JdR. Millán, Closed-loop electrical neurostimulation: Challenges and opportunities, Curr. Opin. Biomed. Eng. 8 (2018) 28–37. doi:10.1016/j.cobme.2018.09.007.

